# Release of stem cells from quiescence reveals multiple gliogenic domains in the adult brain

**DOI:** 10.1101/738013

**Authors:** Ana C. Delgado, Angel R. Maldonado-Soto, Violeta Silva-Vargas, Dogukan Mizrak, Thomas von Känel, Alex Paul, Aviv Madar, Henar Cuervo, Jan Kitajewski, Chyuan-Sheng Lin, Fiona Doetsch

## Abstract

Quiescent neural stem cells (NSCs) in the adult ventricular-subventricular zone (V-SVZ) have a regional identity and undergo activation to generate neurons. The domains for gliogenesis are less explored. Here we show that Platelet-Derived Growth Factor Receptor beta (PDGFRβ) is expressed by adult V-SVZ NSCs that generate olfactory bulb interneurons and glia with slow baseline kinetics. Selective deletion of PDGFRβ in adult V-SVZ NSCs leads to their release from quiescence uncovering multiple domains in the septal wall for oligodendrocyte and astrocyte formation. Unexpectedly, we identify a novel intraventricular oligodendrocyte progenitor inside the brain ventricles. Together our findings reveal different NSC spatial domains for gliogenesis in the adult V-SVZ that are largely quiescent under homeostasis and may have key functions for brain plasticity.

---

Stem cells in the ventricular-subventricular zone (V-SVZ) generate thousands of neurons each day that migrate to the olfactory bulb. Adult NSCs have a regional identity and depending on their spatial location in the V-SVZ give rise to different subtypes of olfactory bulb interneurons (*1*). The adult V-SVZ also generates low numbers of glia under baseline conditions, but whether multiple gliogenic domains exist in the V-SVZ is unknown.

The V-SVZ extends along the lateral ventricles, along both the lateral wall, adjacent to the striatum, and the septal wall, adjacent to the septum, which is far less studied. V-SVZ NSCs are largely quiescent, and both intrinsic and extrinsic signals actively maintain this state. To uncover potential regulators of adult V-SVZ NSC quiescence, we compared the transcriptomes of purified quiescent and activated adult V-SVZ NSCs (*2*), and identified Platelet-Derived Growth Factor Receptor beta (PDGFRβ), a tyrosine kinase receptor, as highly enriched in quiescent adult V-SVZ NSCs (qNSCs).

To characterize PDGFRβ expression *in vivo* in the V-SVZ, we performed immunostaining with lineage markers (Fig. 1A, D and Fig. S1A-H). Adult NSCs are radial GFAP^+^ cells (*2-4*) that contact the lateral ventricle at the center of pinwheels formed by ependymal cells (*3*) and have a primary cilium (*5*). In whole mount preparations, PDGFRβ was expressed by NSCs at the center of pinwheels throughout the rostral-caudal extent of the lateral ventricle (Fig. 1A, C and Fig. S1B-C), with their typical radial morphology revealed by electroporation of a PDGFRβ::mCherry reporter plasmid (Fig. 1B). PDGFRβ was enriched at the apical domain of qNSCs touching the ventricle (Fig. 1D and Fig. S1E-F). As NSCs become activated, they upregulate epidermal growth factor receptor (EGFR). PDGFRβ was also expressed in some activated NSCs (aNSCs, GFAP^+^ EGFR^+^), which had low EGFR expression (Fig. 1D and Fig. S1G), but not in transit amplifying cells (TACs, GFAP^-^ EGFR^+^) (Fig. 1D and Fig. S1G) or doublecortin (DCX^+^) neuroblasts (Fig. S1H). As previously reported (*6*), PDGFRβ was also expressed in CD13^+^ pericytes throughout the brain (Fig. S1D). Therefore within the V-SVZ lineage, PDGFRβ is expressed in adult V-SVZ NSCs, which are largely quiescent.

**Fig. 1.**
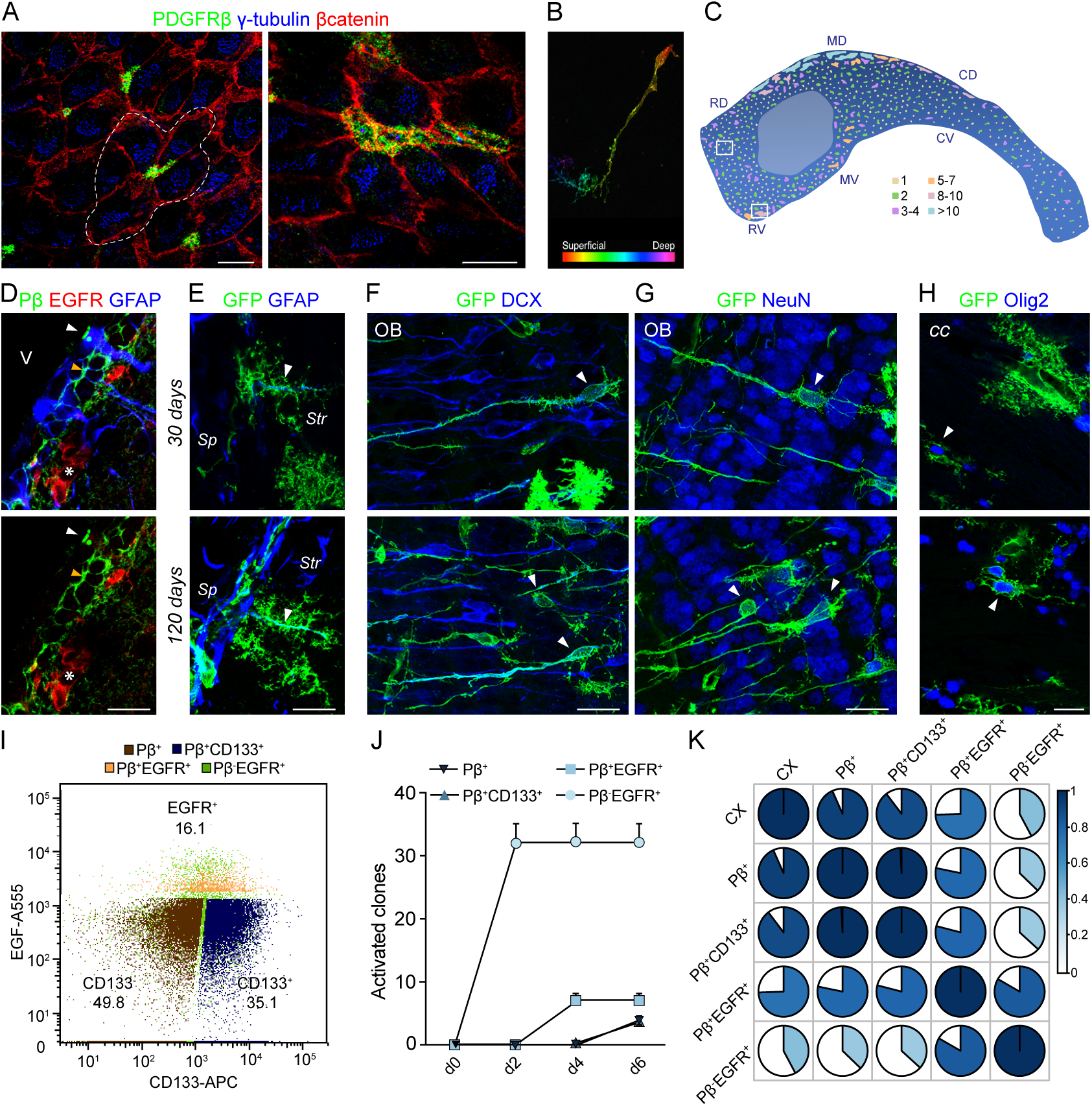
PDGFRβ is expressed in adult V-SVZ NSCs. (A) Confocal images of whole mounts immunostained for β-catenin (red) and γ-tubulin (blue) showing PDGFRβ^+^ NSCs (green) at the center of ependymal cell pinwheels in the rostro-dorsal V-SVZ (one pinwheel is outlined by dashed line) (left). Right shows large cluster of PDGFRβ^+^ NSCs in the rostral-ventral V-SVZ. (B) Confocal Z-stack projection of whole mount showing radial morphology of PDGFR::mCherry electroporated cell. Color code indicates depth of cell relative to the ventricular surface. (C) Map of surface of ventricular wall showing distribution and size of clusters of PDGFRβ^+^ NSCs throughout rostro-caudal extent of the lateral wall. Color-code indicates number of NSCs in each cluster; rostro-dorsal (RD), rostral-ventral (RV), mid-dorsal (MD), mid-ventral (MD), caudal-dorsal (CD), caudal-ventral (CV). Boxes in (C) show location of images in A. (D) Coronal section immunostained for PDGFRβ (green), GFAP^+^ (blue) and EGFR^+^ (red). PDGFRβ is expressed in both quiescent GFAP^+^ (orange arrowhead) and some activated GFAP^+^ EGFR^+^ (white arrowhead) NSCs, but not in GFAP^-^ EGFR^+^ TACs (asterisks). (E-H) Lineage-tracing of PDGFRβ+ cells. Coronal section showing GFP^+^ (green) GFAP^+^ (blue) cells with radial morphology in V-SVZ at 30 and 120 days (white arrowheads). (F, G) Coronal sections of olfactory bulb (OB) showing GFP^+^ DCX^+^ neuroblasts (F) and GFP^+^ NeuN^+^ mature neurons (G) (white arrowheads) in the granule cell layer at 30 and 120 days. (H) GFP^+^ (green) Olig2^+^ (blue) oligodendrocytes in the *cc* at 30 and 120 days (white arrowheads). (I) Representative FACS multigraph overlay showing distribution of PDGFRβ^+^ cells in GFAP::GFP^+^CD24^-^ subpopulations gated on EGFR and CD133. PDGFRβ^+^ cells are shown in dark blue, brown or orange. (J) Activation kinetics of FACS purified populations in vitro. (K) Correlation of genome-wide gene expression profiles across different populations. V, ventricle; *Str*, striatum; *Sp*, septum; *cc*, corpus callosum. Scale bars, A, D-H 10µm.

PDGFRβ^+^ V-SVZ cells gave rise to neuronal and glial progeny and were long-term neurogenic and gliogenic *in vivo*, as shown by lineage tracing using PDGFRβ-P2A*-* CreER^T2^;mT/mG mice (*7*) (Fig. S1I). We detected GFP^+^ immature Dcx^+^ and mature NeuN^+^ neurons in the olfactory bulb, as well as GFP^+^ Olig2^+^ oligodendrocytes in the corpus callosum 30 days after tamoxifen injection, and their numbers increased over time (Fig. 1F-H). GFP^+^ GFAP^+^ radial cells (Fig. 1E), EGFR^+^ transit amplifying cells (Fig. S1J) and migrating neuroblasts (Fig. S1K) were still present in the V-SVZ at both 30 and 120 days, revealing that PDGFRβ^+^ V-SVZ stem cells continue to generate progeny four months after induction. In addition, both GFP^+^ pericytes and stellate astrocytes scattered throughout the brain parenchyma (data not shown) were present consistent with *pdgfrb* expression in some parenchymal astrocytes (*8*); data not shown).

To assess the molecular and functional properties of PDGFRβ^+^ V-SVZ NSCs, we FACS purified them from GFAP::GFP mice. The vast majority of GFP^+^ PDGFRβ^+^ cells were EGFR negative, and corresponded to our previously described CD133^+^ and CD133^-^ GFP^+^ populations (PDGFRβ^+^ CD133^+^ GFAP::GFP^+^ (Pβ^+^ CD133^+^) and PDGFRβ^+^ GFAP::GFP^+^(Pβ^+^), respectively) (Fig. 1I) (*2*). Interestingly, PDGFRβ resolved the EGFR^+^ aNSCs into two populations, one of which was PDGFRβ^+^ (PDGFRβ^+^ EGFR^+^ GFAP::GFP^+^ (Pβ^+^ EGFR^+^)) and the other negative (PDGFRβ^-^ EGFR^+^ GFAP::GFP^+^ (Pβ^-^EGFR^+^)) (Fig. 1I).

RNA sequencing and functional analysis of the FACS purified populations revealed that expression of PDGFRβ was inversely correlated with EGFR expression and cell division (Fig. S1L). The kinetics of colony formation reflected expression of PDGFRβ, with Pβ^+^ EGFR^-^ populations having slower kinetics than EGFR^+^ populations, and the Pβ^+^ EGFR^+^ population in between (Fig. 1J and Fig. S1M). Consistent with this, genome-wide correlation analysis of the FACS purified populations showed that Pβ^+^ EGFR^+^ cells had correlation scores intermediate between Pβ^+^ only and Pβ^-^ EGFR^+^ populations (Fig. 1K), whereas Pβ^+^ only populations were similar to each other and to cortical astrocytes. While both Pβ^+^ CD133^+^ and Pβ^+^ populations contain qNSCs, the Pβ^+^ CD133^+^ population is better defined (*2*). We therefore focused on Pβ^+^ CD133^+^ qNSCs for molecular comparisons (Fig. S1N-Q). The top GO pathways enriched in Pβ^+^ CD133^+^ qNSCs versus the Pβ^+^ EGFR^+^ population highlighted pathways implicated in quiescence such as the regulation of lipid metabolism, G-protein signaling, cell adhesion, and immune response (Fig. S1N) (*2, 9*), as well as novel pathways, including sirtuin 6 regulation, circadian rhythm, S1PR1 signaling and PDGF signaling (Fig. S1N). In contrast, Pβ^+^ EGFR^+^ cells as compared to qNSCs were enriched in pathways related to entry into cell cycle, regulation of translation initiation, Notch signaling, and oligodendrocyte differentiation from adult NSCs (Fig. S1O). Consistent with their faster *in vitro* activation kinetics, the majority of pathways enriched in Pβ^-^ EGFR^+^ versus Pβ^+^ EGFR^+^ cells were cell-cycle and cytoskeletal remodeling related (Fig. 1Q). Together these data show that PDGFRβ expression is linked to the quiescent state, and suggest it must be down-regulated for proliferation.

To functionally assess the effect of deleting PDGFRβ selectively in adult GFAP^+^ V-SVZ NSCs *in vivo*, we generated inducible triple transgenic hGFAP::CreER^T2^;PDGFRβ^+/+^;A14 (PDGFRβ^WT^) and mutant hGFAP::CreER^T2^;PDGFRβ^fl/fl^;A14 (PDGFRβΔ) mice in which CreER^T2^ recombinase is under the control of the human GFAP promoter (hGFAP::CreER^T2^; (*10*). Upon tamoxifen injection, the first and second immunoglobulin domains of the PDGFRβ gene are deleted preventing PDGF-ligand binding (*11-13*), and tdTomato is expressed allowing cell fate to be followed (Fig. 2A). Deletion of PDGFRβ led to an increase in activated stem cells (Tom^+^ GFAP^+^ EGFR^+^) and dividing stem cells (Tom^+^ GFAP^+^ Ki67^+^ cells) (Fig. 2B-E) one day after ending tamoxifen injections (1dpi). Importantly, increased activation occurred in both the lateral and septal walls. TACs (Tom^+^ GFAP^-^ EGFR^+^) and neuroblasts (Tom^+^ DCX^+^) were also increased (Fig. 2B and Fig. S2B-C). We also detected a small increase in Tom^+^ Caspase3^+^ cells in PDGFRβΔ mice, consistent with a role for PDGFRβ in cell survival in other cell types (*14-16*) (Fig. S2D-E). However, the proportion of dying Tom^+^ cells was very low compared to the large increase in total Tom^+^ cell number (Fig. S2A). Thus, in PDGFRβΔ mice, stem cells in both the lateral wall and septal wall are released from quiescence. This increased activation of NSCs in PDGFRβΔ mice resulted in more mature Tom^+^ NeuN^+^ neurons in the olfactory bulb (Fig. 2F, G), and more Tom^+^ Olig2^+^ oligodendrocytes in the corpus callosum than in control mice at 45 or 180 dpi (Fig. 2H, I). Stem cell activation and progeny were also still increased at later timepoints in the V-SVZ (Fig. S2F-I). Importantly, more qNSCs also became activated *in vitro*, with faster activation kinetics, when FACS purified qNSCs were cultured with PDGFDD ligand and PDGFRβ blocking antibodies, but not with isotype controls (Fig. 2J, K).

**Fig. 2.**
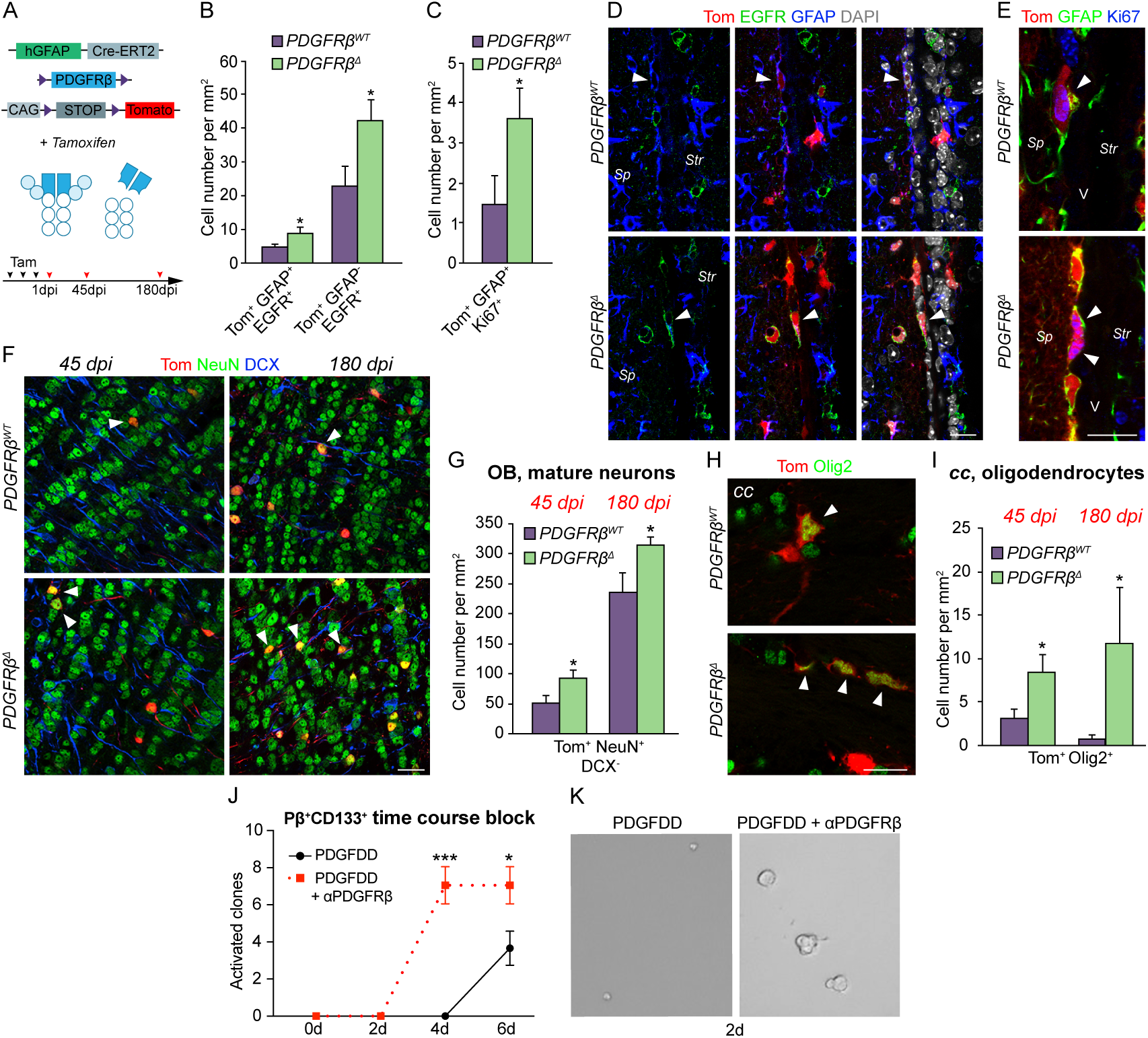
PDGFRβ deletion releases adult NSCs from quiescence. (A) Schema of experimental paradigm. Upon recombination the first and second immunoglobulin domains of PDGFRβ are deleted. Mice were injected with tamoxifen for 3 days and sacrificed 1, 45 or 180 dpi later. (B, C) Quantification of (B) Tom^+^ aNSCs (Tom^+^ GFAP^+^ EGFR^+^), TACs (Tom^+^ GFAP^-^ EGFR^+^) and (C) Tom^+^ GFAP^+^ dividing cells (Tom^+^ GFAP^+^ Ki67^+^) in V-SVZ of PDGFRβ^WT^ and PDGFRβΔ mice. (D, E) PDGFRβ^WT^ and PDGFRβΔ V-SVZ coronal sections immunostained for (D) Tom (red), EGFR (green) and GFAP (blue) or (E) Tom (red), GFAP (green) and Ki67 (blue) and DAPI (white). White arrowheads indicate Tom^+^ aNSCs (D) and dividing GFAP^+^ cells (E). (F) Coronal section of granule cell layer in OB immunostained for Tom (red), NeuN (green) and DCX (blue) at 45 and 180 dpi. (G) Quantification of mature neurons (Tom^+^ NeuN^+^ DCX^-^) in OB in PDGFRβ^WT^ vs PDGFRβΔ mice. (H) Confocal images of Tom^+^ (red) Olig2^+^ (green) cells (white arrowheads) in the corpus callosum (cc) of PDGFRβ^WT^ and PDGFRβΔ mice. (I) Quantification of oligodendrocytes (Tom^+^ Olig2^+^) in the *cc*. (J) Time course of appearance of activated clones from Pβ^+^ CD133^+^ qNSCs treated with αPDGFRβ antibodies in presence of PDGFDD. (K) Images of qNSCs treated with PDGFDD (left) or PDGFDD and αPDGFRβ antibodies (right), which promotes activation. *Str*, striatum; *Sp*, septum. Scale bars, D-F, H 10um. *p<0.05, ***p<0.001 Error bars indicate SEM.

PDGFRβ is therefore expressed in qNSCs in the adult V-SVZ, and is down-regulated as stem cells activate. Indeed, deletion of PDGFRβ selectively in adult NSCs releases them from quiescence, in contrast to deleting PDGFRβ embryonically (*17, 18*). As such, PDGFRβ likely exerts distinct functions at different stages of brain development (*17, 19-21*) and in different cell types, such as pericytes (*22, 23*).

The release of adult NSCs from quiescence in PDGFRβΔ mice provides a powerful tool to uncover V-SVZ NSC domains *in vivo* that are normally more quiescent. In the adult V-SVZ, olfactory bulb interneurons arise from throughout the lateral wall and rostral levels of the septal wall, and oligodendrocytes from the dorsal-lateral V-SVZ (*1, 24, 25*). Little is known about astrocyte formation. Intriguingly, PDGFRβΔ mice revealed several novel gliogenic domains in the adult V-SVZ, especially in the septal wall.

We identified several GFAP^+^ cell types with different distributions and morphologies in the V-SVZ. Previously described radial cells with a process perpendicular to the ventricle were present in both walls, but more enriched in the dorsal and ventral V-SVZ (Fig. 3A, B and Fig. S3A). A second radial cell type with a process parallel to the ventricle was more prevalent on the septal side (Fig. 3A, B and Fig. S3B). We also identified cells with a rounded, plump soma with short small GFAP+ processes (Fig. 3A, B and Fig. S3C), enriched in the septal side and rare under baseline conditions, which we named “gorditas”. Finally, mature highly branched (stellate) astrocytes, with low GFAP expression were also present (Fig. S3D). All GFAP^+^ cell types expressed glutamine synthetase (GS). The release from quiescence did not affect all GFAP+ populations equally in PDGFRβΔ mice. Mature stellate astrocytes were unchanged in the V-SVZ and in the cortex at 1dpi (Fig. S3H). In contrast, radial cells significantly increased in both the lateral and septal V-SVZ (Fig. 3C), consistent with symmetric self-renewing division of NSCs (*26*). Interestingly, gorditas were only significantly increased in the septal V-SVZ of PDGFRβΔ mice, largely in the mid-to-ventral area (Fig. 3A-C), suggesting that this region is a domain for astrocyte generation. Indeed, stellate Tom^+^ GS^+^ GFAP^-^ cells were increased in the septum at early timepoints (Fig. S3E, F). Proliferating cells were only localized near the V-SVZ (Fig. 3D and Fig. S3G) and we did not detect Tom^+^ MCM2^+^ or BrdU^+^ cells in the septum (Fig. 3D and Fig. S3G), suggesting that parenchymal proliferation does not account for this increase. At 45dpi, Tom^+^ stellate astrocytes were greatly increased in the septum itself (Fig. 3E, G), away from the V-SVZ, and the increase in Tom^+^ gorditas in the septal V-SVZ was maintained (Fig. 3E-F). This increase in stellate astrocytes was specific to the septum, indicating that this brain region may have unique requirements for astrocytes.

**Fig. 3.**
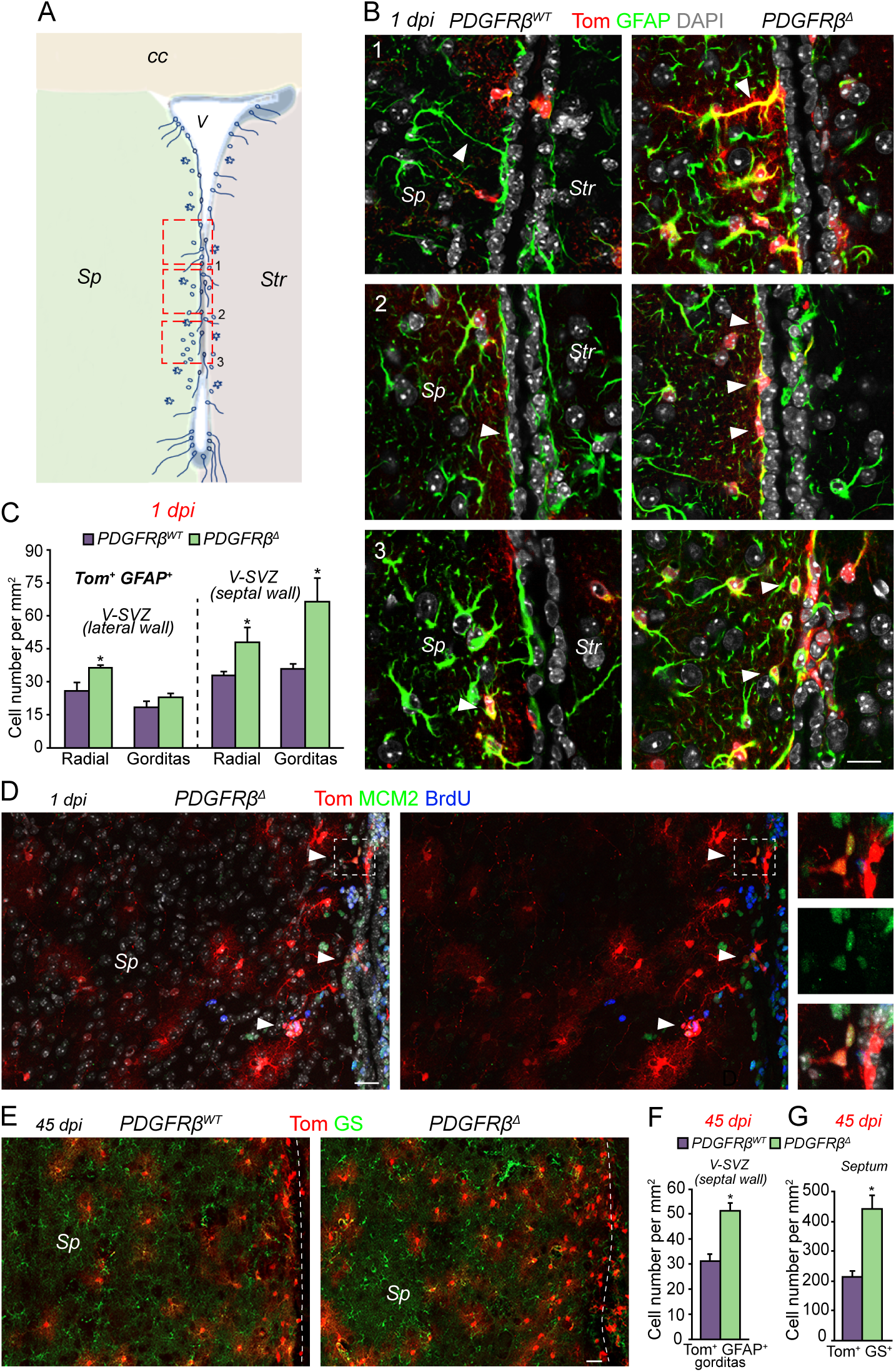
Novel astrogenic domain in the adult septal V-SVZ. (A) Coronal schema depicting different types and distribution of GFAP^+^ cells in adult V-SVZ. Boxes and numbers show location of images shown in B. (B) Confocal images showing different types of Tom^+^ GFAP^+^ cells at 1dpi in PDGFRβ^WT^ (left) and PDGFRβΔ (right) mice. 1, Radial cells perpendicular to the ventricle (white arrowhead). 2, Radial cells parallel to the ventricle (white arrowheads). 3, Gorditas were predominantly located in the septal V-SVZ (white arrowheads). (C) Quantification of GFAP^+^ radial cells and gorditas in the lateral and septal V-SVZ. (D) Coronal section of the septal wall and part of the septum immunostained for Tom (red), BrdU (blue) and MCM2 (green). Tom+ proliferating cells (white arrowheads) are close to the ventricle at 1dpi in PDGFRβΔ mice. Right panels show higher power magnification of proliferating Tom^+^ cells in area in box. (E) Images of Tom^+^ (red) and Glutamine Synthetase (GS, green) immunostaining in septal wall and septum at 45 dpi showing large increase in Tom^+^ GS^+^ cells in PDGFRβΔ as compared to PDGFRβ^WT^ mice. Dashed line indicates ventricle. (F, G) Quantification of (F) Tom^+^ gorditas in septal V-SVZ and (G) Tom^+^ GS^+^ stellate astrocytes in the septum at 45 dpi. *Str*, striatum; *Sp*, septum; *cc*, corpus callosum. Scale bars, B 10um; D, E 20um. *p<0.05. Error bars indicate SEM.

The release from quiescence revealed that the septal V-SVZ also contains oligodendrogenic domains. A restricted region containing Tom^+^ Olig2^+^ cells and many BrdU^+^ cells was turned on in the dorso-medial corner of the septal V-SVZ near the corpus callosum in PDGFRβΔ mice (Fig. 4A, B, D, E). Scattered Tom^+^ Olig2^+^ cells were also present along the dorso-ventral length of the septal V-SVZ. In addition, Tom^+^ Olig2^+^ cells were greatly increased in the corpus callosum at 1dpi (Fig. 4D and Fig. S4A). The septal wall therefore contains several quiescent NSC domains that are gliogenic.

**Fig. 4.**
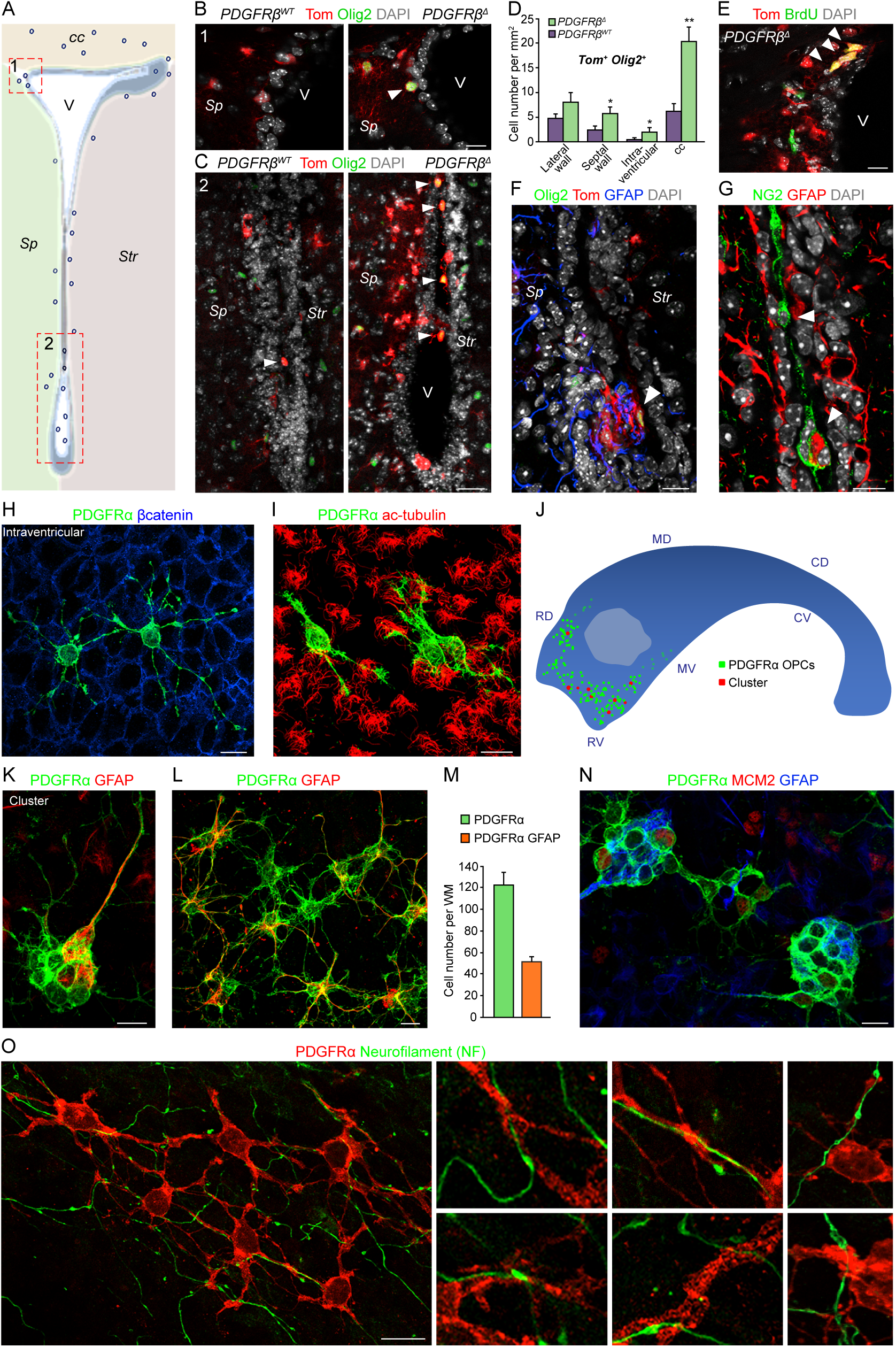
Novel oligodendrogenic domains in the septal V-SVZ and inside the ventricles. (A) Coronal schema of V-SVZ showing location of OPC domains revealed by PDGFRβΔ mice. Boxes and numbers show location of images in B and C. (B) Image of Tom^+^ (red) and Olig2^+^ (green) cells (white arrowheads) in dorsal-medial corner of V-SVZ in PDGFRβ^WT^ and PDGFRβΔ mice. (C) Images of ventral V-SVZ showing large numbers of intraventricular Tom^+^ (red) Olig2^+^ (green) cells (white arrowheads) in PDGFRβΔ as compared to PDGFRβ^WT^ mice. (D) Quantification of Tom^+^ Olig2^+^ cells in different domains in PDGFRβ^WT^ and PDGFRβΔ mice. (E) Dorsal-medial corner of V-SVZ shows an increase of Tom^+^ (red) BrdU^+^ (green) cells (white arrowheads) in PDGFRβΔ mice. (F) Intraventricular Tom^+^ (red) GFAP^+^ (blue) cluster with Olig2^+^ (green) OPCs (white arrowheads) in PDGFRβΔ mice. (G) Intraventricular GFAP^+^ (red) /NG2^+^ (green) cluster (lower arrowhead) with several nearby individual NG2^+^ OPCs (upper arrowhead) in wild type mice. (H) Confocal image of whole mount from wild type mouse showing PDGFRα^+^ OPCs on the surface of ependymal cells, visualized by β-catenin immunostaining. (I) Confocal image of whole mount from wild type mouse showing PDGFRα^+^ OPCs between the cilia of ependymal cells (red, acetylated tubulin). (J) Map showing distribution of intraventricular OPCs and clusters with GFAP+ cells. (K, L) Confocal images of whole mounts showing (K) GFAP+ radial cell with a PDGFRα^+^ cluster and (L) individual PDGFRα^+^GFAP^+^ and PDGFRα^+^ only cells. (M) Quantification of PDGFRα^+^ only and PDGFRα^+^GFAP^+^ cells per whole mount. (N) Whole mount immunostained for PDGFRα^+^ (green), GFAP (blue) and MCM2 (red) showing some dividing OPCs. (O) Whole mount immunostained for PDGFRα^+^ (red) and neurofilament (NF, green) showing association of intraventricular OPCs with supraependymal axons. Panels at right show higher magnification images. *Sp*, septum; *Str*, striatum; *cc*, corpus callosum; V, ventricle. Scale bars, 10µm. *p<0.05, **p<0.01. Error bars indicate SEM.

Strikingly, we also identified a novel oligodendrocyte progenitor (OPC) type inside the brain ventricles. In PDGFRβΔ mice, intraventricular Tom^+^ Olig2^+^ cells derived from GFAP^+^ NSCs were attached to the luminal surface of the wall (Fig. 4A, C) close to large clusters of Tom^+^ GFAP^+^ cells (Fig. 4F). Intraventricular OPCs expressed typical OPC markers (Olig2^+^, NG2^+^ and PDGFRα^+^) and, importantly, were also present in wild type brains, closely associated with intraventricular clusters of GFAP^+^ and NG2^+^ cells (Fig. 4G). To clearly visualize the distribution and morphology of intraventricular OPCs in wild type mice, we used whole mount preparations of the surface of the wall. Intraventricular OPCs were localized on top of ependymal cells (Fig. 4H and Fig. S4B) nestled between their cilia (Fig. 4I and Fig. S4E) in the rostral portion of the anterior horn of the ventricle (Fig. 4J), and frequently contacted other PDGFRα^+^ cells (Fig. 4H, O, and Fig. S4I, K, L). Their morphology was distinct from PDGFRα^+^ OPCs in the brain parenchyma (Fig. S4B, C). Supporting their origin from GFAP+ cells, intraventricular clusters containing radial GFAP^+^ cells, most of which co-expressed PDGFRα, were tightly associated with PDGFRα^+^ only cells (Fig. 4K, N and Fig. S4D). Nearby, resolving clusters of dispersed PDGFRα^+^ GFAP^+^ double-labeled cells and PDGFRα only cells were present. Some, but not all PDGFRα^+^ cells were dividing based on MCM2 and Ki67 expression (Fig. 4M, N and Fig. S4G, H). Interestingly, many intraventricular OPCs were closely apposed to or partially enwrapped supraependymal axons from neurons in other brain regions present on the surface of the walls of the lateral ventricles (Fig. 4O and Fig. S4L). These neurons regulate both NSC proliferation (*27*) and ependymal cilia beating (*28*). Importantly, we did not detect mature myelinating Rip, MOG or MBP^+^ oligodendrocytes inside the ventricles (Fig S4I to K), suggesting that intraventricular OPCs have a unique function.

In sum, we uncover multiple V-SVZ domains for the generation of glia (Fig. S4M) and that the septal V-SVZ contains a reservoir of quiescent NSCs for the generation of astrocytes and oligodendrocytes (*29*), suggesting that the septum may have specialized requirements for glial cells that may be recruited on demand in different contexts. Furthermore, the identification of intraventricular OPCs that are bathed by cerebrospinal fluid and contact axons from distant brain regions, and are therefore poised to dynamically sense signals in the CSF and from other brain areas, further highlights glial diversity in the adult brain and opens new vistas for understanding the role of these new types of glia in health and disease.

## Acknowledgements

We thank members of the Doetsch and Wichterle labs and D. Thaler for discussion; V. Crotet for genotyping; the Biozentrum Imaging Core Facility (IMCF); K. Gordon and S. Tetteh of the Herbert Irving Comprehensive Cancer Center Flow Cytometry Core of Columbia University for assistance with FACS; and J. Bögli at the Biozentrum FACS Core Facility.

## Author Contributions

Conceptualization: F.D., A.R.M-S., A.C.D.; Performed experiments: A.C.D., A.R.M-S., V.S-V., D.M., T.v.K., A.P.; Data Analysis: A.C.D., A.R.M-S., V.S-V., D.M., A.M.; Supervision: F.D.; Funding acquisition: F.D.; Manuscript writing: A.C.D., A.R.M-S., V.S-V., D.M. and F.D.; Resources: H.C., J.K. and C-S.L contributed PDGFRβ-P2A-CreER^T2^;mT/mG mice.

## Data and materials availability

All data will be deposited in the National Center for Biotechnology Information Gene Expression Omnibus. Accession numbers have been requested.

