## Supplementary Materials for "Release of stem cells from quiescence reveals multiple gliogenic domains in the adult brain"

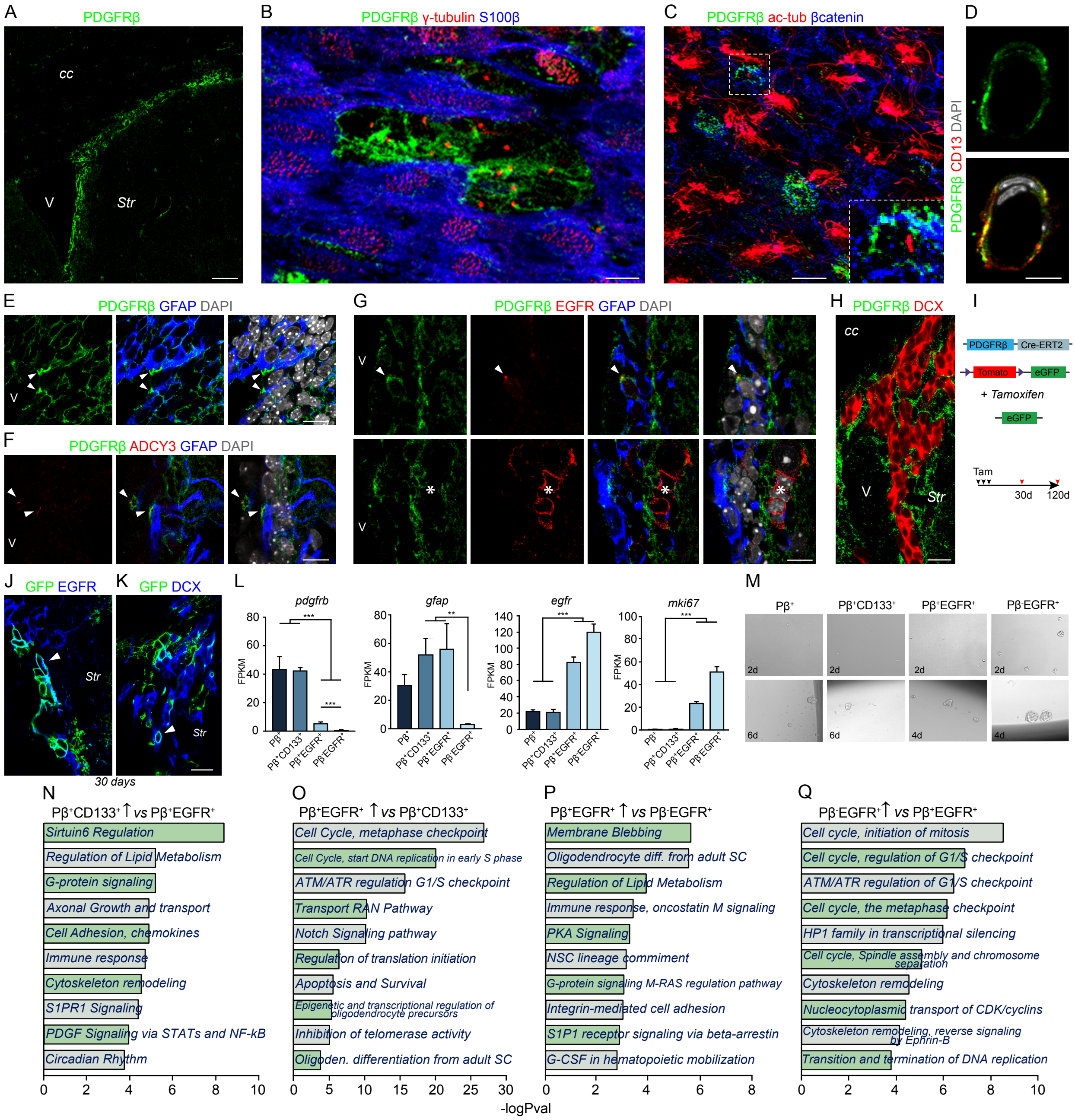


**Fig. S1 Functional and molecular properties of PDGFRβ+ NSCs in the adult V-SVZ**

(A) Coronal section showing high levels of PDGFRβ (green) in the V-SVZ and dorsal lateral aspect of the V-SVZ. Labeled cells throughout the tissue are pericytes (see S1D). (B) Confocal image of whole mount showing large cluster of PDGFRβ$^+$ γ-tubulin$^+$ NSCs (green) (red) surrounded by S100β$^+$ (blue) PDGFRβ-negative ependymal cells in the mid-dorsal area. (C) Confocal image of whole mount showing PDGFRβ$^+$ NSCs (green) with a primary cilium labeled by acetylated tubulin (red). Inset shows higher magnification view of region in box. (D) PDGFRβ (green) labels CD13$^+$ pericytes (red) throughout the brain. Nuclei are labeled with DAPI. (E) Confocal images of coronal section showing apical enrichment of PDGFRβ in ventricle-contacting GFAP$^+$ (blue) NSCs. DAPI is in grey. (F) Confocal images of coronal section showing apical enrichment of PDGFRβ in ventricle-contacting $GFAP^+$ (blue) NSCs with a primary cilium (adenylate cyclase (ADCY3), red). DAPI is in grey. (G) Confocal images of coronal sections of V-SVZ showing that PDGFRβ (green) is expressed in $EGFR^+$ low (red) $GFAP^+$ (blue) NSCs (arrowhead) (upper panels), but not in $GFAP^-$ cells with high levels of $EGFR^+$ expression (asterisks) (lower panels). (H) Confocal image of coronal section showing that $doublecourtin^+$ (DCX, red) neuroblasts do not express PDGFRβ (green). (I-K) Lineage tracing of $PDGFR\beta^+$ cells using PDGFRβ-P2A $CreER^{T2}$; mT/mG mice. (I) Schema of mice and experimental paradigm. PDGFRβ-P2A $CreER^{T2}$; mT/mG mice were injected for 3 days with tamoxifen and sacrificed 30 or 120 days later. (J) Images of $GFP^+$ $EGFR^+$ transit amplifying cells in the V-SVZ (white arrowhead) at 30dpi. (K) Images of $GFP^+$ $DCX^+$ neuroblasts (white arrowhead) in the V-SVZ at 30 dpi. (L) Expression of selected genes (FPKM) in purified populations. Asterisks indicate significant differential expression (fdr < 0.05). (M) Images of activated clones from FACS purified populations at different time points. (N-Q) Selected top-enriched pathway maps from Metacore analysis comparing $P\beta^+CD133^+$ qNSCs to $P\beta^+$ $EGFR^+$ (N), $P\beta^+$ $EGFR^+$ to $P\beta^+CD133^+$ qNSCs (O), $P\beta^+$ $EGFR^+$ to $P\beta^-$ $EGFR^+$ (P) and $P\beta^-$ $EGFR^+$ to $P\beta^+$ $EGFR^+$ (Q) populations. Pathways are ranked by -log(p-val). $^{**}p<0.01$, $^{***}p<0.001$. Error bars indicate SEM. V, ventricle; *Str*, striatum; *cc*, corpus callosum. Scale bars, A 50um; B, C 5um; D-H, J, K 10um.

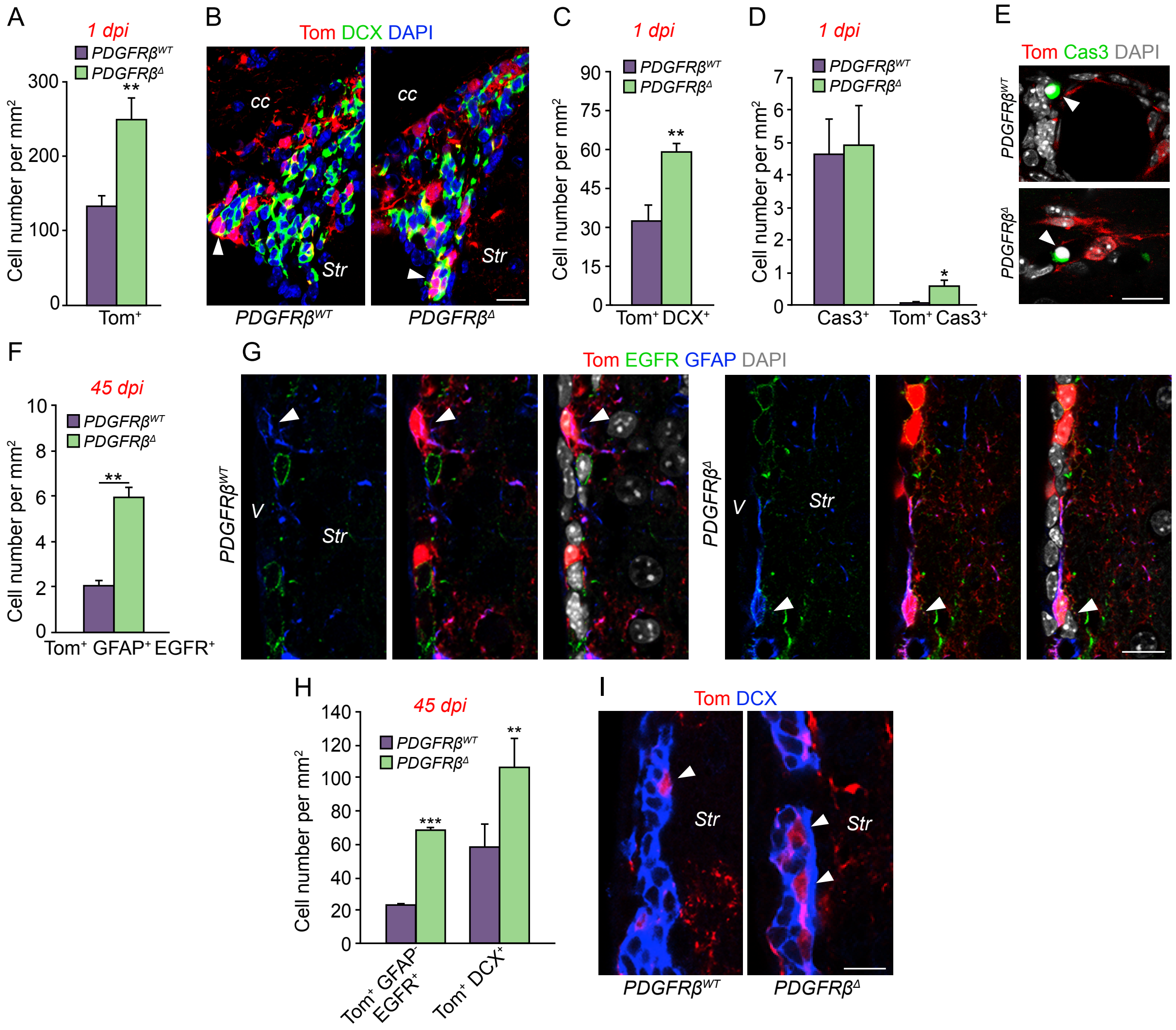


**Fig. S2. PDGFRβ deletion affects NSC proliferation and cell survival**

(A) Quantification of total Tom$^{+}$ cells in PDGFRβ$^{WT}$ and PDGFRβ$^{\Delta}$ mice. (B) Coronal section immunostained for Tom (red) and DCX (green) and DAPI (blue) in V-SVZ PDGFRβ$^{WT}$ and PDGFRβ$^{\Delta}$ mice. White arrowheads indicate Tom$^{+}$ neuroblasts. (C) Quantification of Tom$^{+}$ neuroblasts (Tom$^{+}$ DCX$^{+}$) in V-SVZ of PDGFRβ$^{WT}$ and PDGFRβ$^{\Delta}$ mice. (D) Quantification of total Caspase3$^{+}$ and Tom$^{+}$ Caspase3$^{+}$ cells in the V-SVZ of PDGFRβ$^{WT}$ and PDGFRβ$^{\Delta}$ mice. (E) Images of Tom$^{+}$ (red) and Caspase3$^{+}$ (green) cells (white arrowheads) in the V-SVZ of PDGFRβ$^{WT}$ and PDGFRβ$^{\Delta}$ mice. (F) Quantification of aNSCs (Tom$^{+}$ GFAP$^{+}$ EGFR$^{+}$) in PDGFRβ$^{WT}$ vs PDGFRβ$^{\Delta}$ mice at 45dpi. (G) V-SVZ coronal sections immunostained for Tom (red), EGFR (green) and GFAP (blue) at 45 dpi in PDGFRβ$^{WT}$ vs PDGFRβ$^{\Delta}$ mice. White arrowheads indicate Tom$^{+}$ aNSCs (GFAP$^{+}$ EGFR$^{+}$). (H) Quantification of TACs (Tom$^{+}$ GFAP$^{-}$ EGFR$^{+}$) and neuroblasts (Tom$^{+}$ DCX$^{+}$) at 45 dpi. (I) Coronal section showing Tom$^{+}$ (red) DCX$^{+}$ (blue) neuroblasts (white arrowheads) at 45dpi. Nuclei were stained with DAPI. Scale bars, B, E, G, I 10µm. *p<0.05, **p<0.01, ***p<0.001. Error bars indicate SEM.

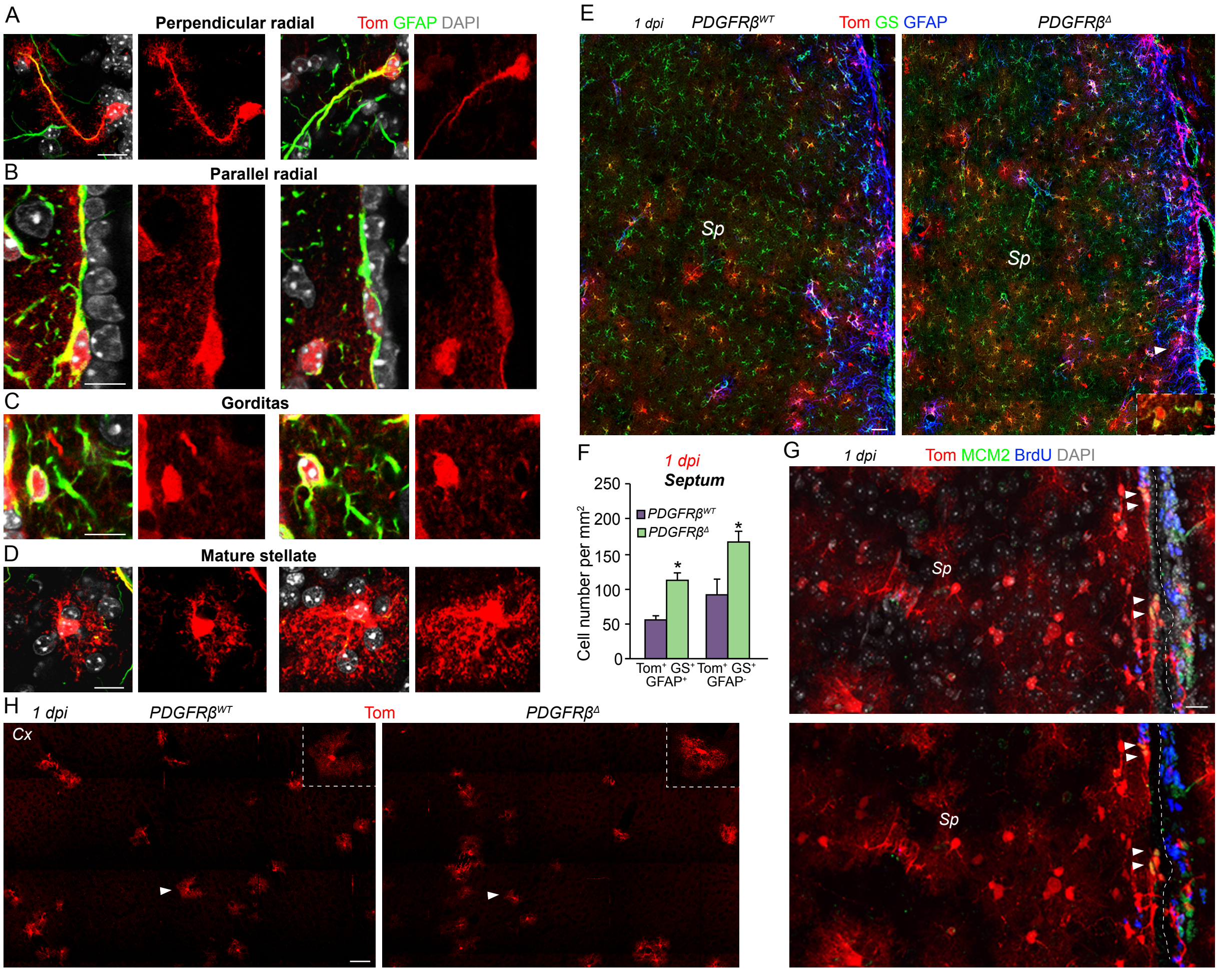


**Fig. S3. Morphologies of GFAP$^+$ cells in the adult V-SVZ and septal wall astrogenic domain**

(A-D) Confocal images showing different morphologies of Tom$^+$ (red) GFAP$^+$ (green) cells at 1dpi. (A) Tom$^+$ GFAP$^+$ radial cells perpendicular to the ventricle. (B) Tom$^+$ GFAP$^+$ radial cells parallel to the ventricle. (C) Tom$^+$ GFAP$^+$ gorditas in the septal wall. (D) Mature highly branched (stellate) Tom$^+$ astrocytes. (E) Tom$^+$ (red) GS$^+$ (green) GFAP$^+$ (blue) cells in septum and septal V-SVZ showing an increase of Tom$^+$ GS$^+$ GFAP$^+$ and Tom$^+$ GS$^+$ GFAP$^-$ cells in PDGFRβ$^{\Delta}$ mice in both regions at 1dpi. Inset shows higher magnification of Tom$^+$ GS$^+$ cells in the septal V-SVZ. (F) Quantification of Tom$^+$ GS$^+$ GFAP$^+$ and Tom$^+$ GS$^+$ GFAP$^-$ cells in septum at 1 dpi. (G) Coronal section of septal V-SVZ and septum immunostained for Tom (red), BrdU (blue) and MCM2 (green). At 1dpi in PDGFRβ$^{\Delta}$ mice proliferation is close to the ventricle (white arrowhead) whereas the septum does not contain Tom$^+$ dividing cells. (H) Images of the cortex showing similar numbers of branched Tom$^+$ astrocytes (white arrowhead) in PDGFRβ$^{WT}$ and PDGFRβ$^{\Delta}$ mice. Inset shows higher magnification. Sp, septum; Cx, cortex. Scale bars, A, D, G 10um; B, C 5um; E, H 20um. *$p<0.05$. Error bars indicate SEM.

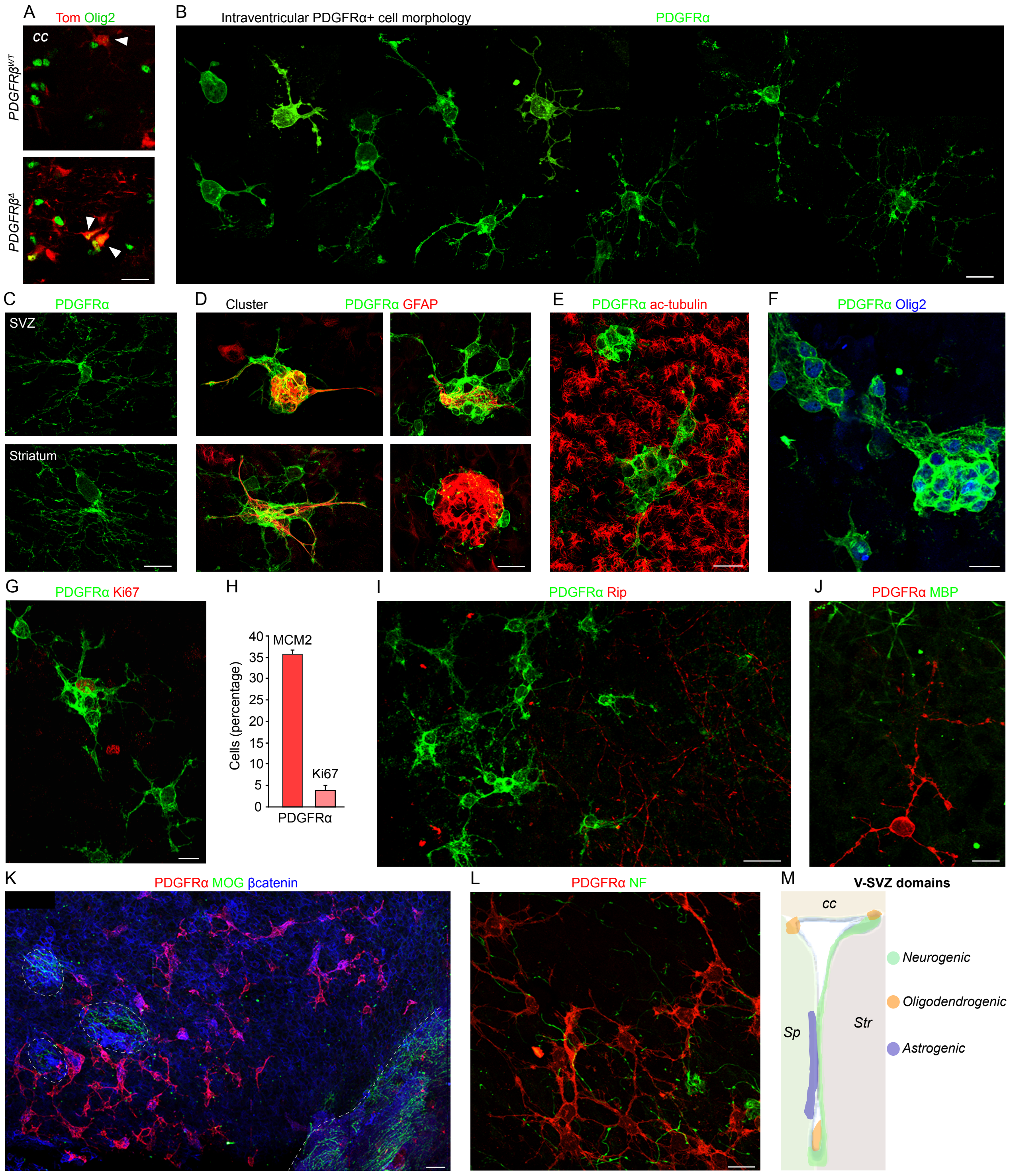


**Fig. S4. Characterization of intraventricular oligodendrocyte progenitors**

(A) Images from coronal sections of $Tom^+$ (red) and $Olig2^+$ (green) cells (white arrowheads) in the corpus callosum of $PDGFR\beta^{WT}$ and $PDGFR\beta^{\Delta}$ mice. (B-G, I-L) Confocal images of intraventricular $PDGFR\alpha^+$ OPCs in whole mount preparations. (B) Collage showing different morphologies of intraventricular $PDGFR\alpha^+$ OPCs from less to more mature. (C) Images showing morphology of parenchymal $PDGFR\alpha^+$ cells deeper in the V-SVZ and in the striatum. (D) Images of intraventricular clusters of PDGFRα$^+$ (green) and GFAP$^+$ (red) cells. (E) Image showing ependymal cell cilia (acetylated tubulin, red) and intraventricular PDGFRα$^+$ OPCs (green). (F) Intraventricular clusters of PDGFRα$^+$ (green) cells express Olig2$^+$ (blue). (G) Image showing that some intraventricular PDGFRα$^+$ (green) OPCs are Ki67$^+$ (red). (H) Quantification of intraventricular PDGFRα$^+$ MCM2$^+$ and PDGFRα$^+$ Ki67$^+$ OPCs. (I) Images of whole mount projection of immunostaining for PDGFRα (green) and Rip (red) showing that PDGFRα$^+$ OPCs are not myelinated. Rip$^+$ processes (red) visible on the right are from deeper levels in the V-SVZ. (J) Whole mount immunostained with PDGFRα (red) and MBP (green) showing that PDGFRα$^+$ OPCs do not express MBP. (K) Panoramic view of whole mount immunostained with PDGFRα (red), MOG (green) and βcatenin (blue) showing that myelinating cells are not present on the surface of the wall, in contrast to the commissure (to right of dashed line). Dashed circles shows myelinated axons in the striatum where the wholemount was nicked during preparation, damaging the ependymal later. (L) Confocal image of PDGFRα (red) and Neurofilament, NF (green) showing association of intraventricular PDGFRα$^+$ OPCs with supraependymal axons. (M) Coronal schema showing V-SVZ domains in the adult enriched for the generation of neurons (green), astrocytes (purple) and oligodendrocytes (orange). Sp, septum, Str, striatum, cc, corpus callosum. Scale bars, A, I, K; 20um, B-G, J, L 10um.

### Materials and Methods

#### Mice

Experiments were performed in accordance with Columbia University IACUC guidelines and approval of the cantonal veterinary office of Basel-Stadt.

All mice used were two to three months old. The following mouse lines were used: CD1 (Charles River), RjOrl:SWISS (Janvier), PDGFRβ-P2A-CreER$^{T2}$;mT/mG mice (*7*), hGFAP::GFP mice (*30*), Jackson Laboratory). We bred hGFAP::CreER$^{T2}$ (*10*), Jackson Laboratory), Pdgfrβ$^{tm11Sor}$(*13*), Jackson Laboratory) and Gt(ROSA)26Sor$^{tm14(CAG\text{-}tdTomato)Hze}$ (Ai14; Jackson Laboratory) mice together to generate triple transgenic mice hGFAP::CreER$^{T2}$; PDGFRβ$^{fl/fl}$; A14 (PDGFRβ$^{\Delta}$ ) or hGFAP::CreER$^{T2}$; PDGFRβ$^{+/+}$; A14 (PDGFRβ$^{WT}$).

#### Tamoxifen and BrdU injections

Cre-mediated recombination in CreER$^{T2}$ transgenic mice was induced by administration of Tamoxifen (Sigma) dissolved at 30 mg/ml in corn oil (Sigma). Mice were injected intraperitoneally with 0.1ml once per day for 3 consecutive days and sacrificed at different points after ending tamoxifen injections: For lineage tracing, mice were analyzed 30 days post injection (dpi) and 120 dpi. For PDGFRβ deletion experiments, mice were sacrificed 1 dpi, 45 dpi or 180 dpi. For BrdU experiments, mice were injected with Tamoxifen for 3 days and with BrdU (50mg/kg, Sigma) 12h before the last Tamoxifen injection and 1h before sacrificing the mice.

#### Cloning

The PDGFRβ promoter was cloned from mouse genomic DNA using the following primers: *AseIF 5'*-GCCATTAATGCTGGGTCAGGCACTCC-3' and *XhoIR 5'-ATACTCGAGCTCCGGGAGGAGCGGAGCA-3'*. The region amplified by these primers contained a CCAAT motif in the promoter necessary to drive a luciferase assay (*31*). In order to insert the promoter into the CherryPicker (Clontech) membrane-bound mCherry expression vector we performed site-directed mutagenesis (sdm) to remove an AgeI site from the promoter. We used the QuickChange Site- Directed Mutagenesis kit (Stratagene) and the following primers: *sdmF 5'-GCACACAGTACCGGCCCTCAGGTCCTCAAAC-3' sdmR 5'-GTTTGAGGACCTGAGGGCCGGTACTGTGTGC*-3'. The promoter with the point mutation was inserted into the CherryPicker vector using *XhoI* and *AgeI* restriction sites and the following primers: *XhoIF 5'-CACCTCGAGACCGCTGGGTCAGGCACTCC-3' AgeIR 5'-ATAACCGGTCTCCGGGAGGAGCGGAGCA-3'*.

**Electroporation**

The electroporation protocol was adapted from (*32*). CD1 mice were deeply anesthetized using Avertin (Sigma). 1 µl of solution containing 5 mg/ml of PDGFRβ::mCherry plasmid in 0.9% saline was injected into the lateral ventricle using the following coordinates relative to Bregma: anterior-posterior, 0.0; lateral, 0.85; ventral, -2.5 mm. Holding the cathode of a square electroporator towards the ipsilateral V-SVZ, five pulses of 200 V (50 ms; separated by 950 ms) were administered and mice sacrificed 7 days later.

**Tissue preparation for Immunohistochemistry**

Mice were anesthetized by intraperitoneal injection with pentobarbital or Avertin and were sacrificed by intracardial perfusion of 4% paraformaldehyde (PFA) in 0.1M phosphate buffer (PB). Brains were extracted from the skull and post-fixed 3 hours for immunostaining for receptors and overnight for the rest of antibodies. Coronal sections were cut at 25µm using a vibratome (Leica VT1000S). Whole mounts were prepared as previously described (*33*).

**Immunostaining**

Whole mounts or sections were incubated in blocking solution (PBS with 2% bovine serum albumin (BSA) and 0.5% Triton-X100 for antibodies against receptors or 0.2-1.5% Triton X-100 for all others) for 60 minutes and then incubated in primary antibodies in blocking solution for 36 hours at 4°C. After washing, sections were incubated with secondary antibodies for 1-2 hours at room temperature. After washing, sections were counterstained with 4',6- Diamidino-2-Phenylindole Dihydrochloride (DAPI, Sigma). Sections were mounted on slides with FluorSave$^{TM}$ (Millipore Corporation).

**Antibodies**

The following primary antibodies were used: anti- Adenylate cyclase (rabbit, 1:100, Santa Cruz Biotechnology); anti-BrdU (rat, 1:400, abcam); anti-CD13 (rat, 1:100, abcam); anti-cleaved caspase 3 (rabbit, 1:100, Cell Signaling); anti-β-Catenin (mouse, 1:200, BD Bioscience); anti-β-Catenin (rabbit, 1:200, Cell signaling); anti-doublecortin, DCX (goat, 1:100, Santa Cruz); anti-DsRed (rabbit, 1:500, Clontech); anti-EGFR (goat,1:100, R&D); anti-EGFR (rabbit,1:100, abcam); anti-GFAP (chicken, 1:600, Millipore); anti-GFP (goat, 1:500, Rockland); anti-GFP (rat, 1:500, Nacalai Tesque, Inc); anti-GLAST (guinea pig, 1:1000, Chemicon); anti-Glutamine synthetase (rabbit, 1:100, abcam); anti-Ki67 (rabbit, 1:100, abcam); anti-MBP (rabbit, 1:100, abcam); anti-MCM2 (rabbit,1:100, Cell signaling); anti-MOG (mouse, 1:100, Millipore) anti-NeuN (mouse, 1:100, Millipore); anti-Neurofilament (chicken, 1:100, abcam); anti-NG2 (rabbit, 1:100, Millipore); anti-O4 (mouse, 1:200, Chemicon); anti-Olig2 (rabbit, 1:100, Millipore); anti-PDGFRβ (rat, 1:50, eBioscience); anti-PDGFRβ (goat, 1:100, R&D); anti-PDGFRα (rat, 1:100, eBioscience); anti-PDGFRα (goat, 1:100, R&D); anti-Rip (mouse, 1:100, DSHB); anti-S100β (rabbit, 1:200, DAKO); anti-Acetylated Tubulin (mouse, 1:1000, Sigma); anti-βIII Tubulin (mouse, 1:500, Covance); anti-γ-tubulin (mouse, 1:200, Sigma). The following secondary antibodies were used: Alexa Fluor-conjugated (488, 647, 568; 1:600, Molecular probes), Cy3-conjugated (1:1000, Jackson ImmunoResearch).

**FACS**

The SVZs were dissected from heterozygous hGFAP::GFP mice (Jackson Labs) or wild-type CD-1 mice (Charles River), digested with papain (Worthington, 1,200 units per 5 mice, 10 min at 37°C) in PIPES solution [120 mM NaCl, 5 mM KCl, 50 mM PIPES (Sigma), 0.6% glucose, 1x Antibiotic/Antimycotic (Gibco), and phenol red (Sigma) in water; pH adjusted to

7.6 and mechanically dissociated to single cells after adding ovomucoid (Worthington, 0.7 mg per 5 mice) and DNAse (Worthington, 1,000 units per 5 mice). Cells were centrifuged for 10 min at 4°C without brake in 22% Percoll (Sigma) to remove myelin. Antibodies against CD13 and CD31 were included to deplete pericytes (CD13) and endothelial cells (CD31). Cell stainings were done in 3 steps: First, cells were incubated for 20 min in anti-biotin PDGFRβ (1:15, eBioscience) or anti-biotin PDGFRβ (1:25, R&D), CD24-A700 (1:150, Biolegend), CD133-APC (clone 13A4, eBioscience), and CD13 BV45 (1:50, BD) and CD31 BV45 (1:50, BD) to deplete perivascular cells. Cells were washed by centrifugation at 1300rpm for 5min. Next, cells were incubated for 10 min with Streptavidin PE-Cy7 (1:500; eBioscience), and washed by centrifugation. Finally, cells were incubated with Alexa Fluor 555- conjugated EGF for 20 min (1:100; Molecular Probes), and washed by centrifugation. All stainings and washes were carried out on ice in 1% BSA, 0.1% Glucose HBSS solution. To assess cell viability, 4',6-diamidino-2-phenylindole (DAPI; 1:1000; Sigma) was added to the cell suspension. All cell populations were isolated in a single sort using a Becton Dickinson FACS Aria II using 13 psi pressure and 100-μm nozzle aperture and were collected in Neurosphere (NS) medium (details below). Gates were set manually using single-color control samples, FMO controls and isotype controls. Data were analyzed with FlowJo 9.3 data analysis software and displayed using bi-exponential scaling.

**RNA-Sequencing**

For RNA-sequencing experiments, RNA was isolated from FACS-sorted cells (three replicates per population) using anti-biotin PDGFRβ (eBioscience). The experiments were performed on the following populations: hGFAP::GFP$^+$ PDGFRβ$^+$ CD133$^-$ EGFR$^-$ CD24$^-$ (Pβ$^+$), hGFAP::GFP$^+$ PDGFRβ$^+$ CD133$^+$ EGFR$^-$ CD24$^-$ (Pβ$^+$ CD133$^+$), hGFAP::GFP$^+$ PDGFRβ$^+$ CD133$^+$ EGFR$^+$ CD24$^-$ (Pβ$^+$ EGFR$^+$) and hGFAP::GFP$^+$ PDGFRβ$^-$ CD133$^+$ EGFR$^+$ CD24$^-$ (Pβ$^-$EGFR$^+$) from the V-SVZ, as well as hGFAP::GFP$^+$ astrocytes from the cortex (CTX) with more than 500 cells per replicate per cell type. Cell populations were directly FACS-sorted into PicoPure RNA isolation kit (Arcturus) and immediately frozen on dry ice to increase RNA yield. All samples were treated with DNAse (Qiagen) during the RNA isolation step. Whole transcriptome amplification was performed using the SMARTer Ultra Low Input RNA Sequencing-HV Kit (Clontech). Sequencing libraries were generated with Nextera DNA preparation kit (Illumina) according to the HV kit's modified Nextera protocol with an input of 5 ng cDNA. Following PCR amplification, library clean-up was performed using Agencourt AMPure XP beads (Beckman Coulter). The resulting libraries were analyzed on an Agilent Bioanalyzer. High quality libraries were sequenced as paired-end 100bp reads on a Illumina HiSeq2000 or 2500 at Columbia Genome Center.

**High Throughput Sequencing Data Analyses**

For RNA-sequencing experiments, an average of 20+ million reads from five cell populations were aligned to the mouse reference genome sequence (mm10) using TopHat v2,1 (*34*) with an average alignment rate of ~80%. Prior to aligning the reads, they were quality-trimmed to remove adapter sequences and ambiguous bases using cutadapt (*35*) and only uniquely mapped concordant mate pairs were used in the downstream analysis. After read mapping, transcripts were assembled using Cufflinks (*34*) and the expression level estimation for each gene was calculated. These values are reported in fragments per kilobase of transcript per million fragments mapped (FPKM). Total read counts for each gene were calculated using HTseq (*36*). The differential expression analyses on the read counts were performed using EdgeR package deposited in Bioconductor (*37*). Adjusted p values were used to determine statistical significance.

***In vitro* experiments**

For all *in vitro* culture experiments, cells were FACS purified as described above using anti-biotin PDGFRβ (R&D).  To assess activation kinetics of activation *in vitro*, FACS purified cells were grown in Neurosphere (NS) medium [DMEM/F12 (Life Technologies) supplemented with 0.6% Glucose (Sigma), 1x HEPES (Life Technologies), 1x Insulin-Selenium-Transferrin (Life Technologies), 1x Antibiotic/Antimycotic (Gibco), N-2 (Life Technologies) and B-27 (Life

Technologies) supplements] with 20 ng/ml EGF (Upstate) and 20 ng/ml bFGF (Sigma) and Heparin 0.7 U /ml (Sigma). Each sorted population was centrifuged after FACS for 10 min at 1300 rpm at 4°C and seeded in NS medium at a density of 1.4 cell/µl in 96 well plates. For quantification of activated clones, clones were quantified at 2 days and later timepoints after plating. Clusters of two or more cells with large, bright, refractant cytoplasm were counted. For PDGFRβ *in vitro* blocking experiments, FACS purified cells were plated with EGF, FGF and Heparin in the presence of rhPDGFDD (20ng/ml; R&D) and anti-PDGFRβ blocking antibodies (3µg/ml; eBioscience) or with isotype control antibodies (3µg/ml; eBioscience). Clones were monitored every two days.

**Image acquisition, quantification and statistical analysis**

Images from immunostained slices were acquired on a LSM 700 or 800 confocal (Zeiss) using 40X and 25X objectives. For *in vivo* quantification, we used at least four sections from rostral to caudal V-SVZ for each animal and counted the entire septal and lateral wall from ventral to dorsal for each section. For quantification in the septum itself, counts were performed from the midline to a distance 30 µm from the ventricle. We quantified cells and measured the area using Fiji software (*38*). Data are presented as average/mm$^2$. Two-way statistical comparisons were conducted by two-tailed unpaired Student's test. Significance was established at *$p<0.05$, **$p<0.01$, ***$p<0.001$. In all graphs, error bars are standard error of the mean (SEM). When comparing more than two datasets, ANOVA analysis was used to determine significance followed by the Bonferroni multiple comparison test on pairwise comparisons using Prism 6 software.
